# Domain Organization of Lentiviral and Betaretroviral Surface Envelope Glycoproteins Modeled with AlphaFold

**DOI:** 10.1101/2021.08.09.455765

**Authors:** Isidro Hötzel

**Author notes:** Corresponding author: Isidro Hötzel, Department of Antibody Engineering, Genentech, South San Francisco, CA 94080.

## Abstract

The surface envelope glycoproteins of non-primate lentiviruses and betaretroviruses share sequence similarity with the inner proximal domain β-sandwich of the human immunodeficiency virus type 1 (HIV-1) gp120 glycoprotein that faces the transmembrane glycoprotein as well as patterns of cysteine and glycosylation site distribution that points to a similar two-domain organization in at least some lentiviruses. Here, high reliability models of the surface glycoproteins obtained with the AlphaFold algorithm are presented for the gp135 glycoprotein of the small ruminant caprine arthritis-encephalitis (CAEV) and visna lentiviruses and the betaretroviruses jaagsiekte sheep retrovirus (JSRV), mouse mammary tumor virus (MMTV) and consensus human endogenous retrovirus type K (HERV-K). The models confirm and extend the inner domain structural conservation in these viruses and identify two outer domains with a putative receptor binding site in the CAEV and visna virus gp135. The location of that site is consistent with patterns of sequence conservation and glycosylation site distribution in gp135. In contrast, a single domain is modeled for the JSRV, MMTV and HERV-K betaretrovirus envelope proteins that is highly conserved structurally in the proximal region and structurally diverse in apical regions likely to interact with cell receptors. The models presented here identify sites in small ruminant lentivirus and betaretrovirus envelope glycoproteins likely to be critical for virus entry and virus neutralization by antibodies and will facilitate their functional and structural characterization.

**Importance:** Structural information on the surface envelope proteins of lentiviruses and related betaretroviruses is critical to understand mechanisms of virus-host interactions. However, experimental determination of these structures has been challenging and only the structure of the human immunodeficiency virus type 1 gp120 has been determined. The advent of the AlphaFold artificial intelligence method for structure prediction allows high-quality modeling of the structures of small ruminant lentiviral and betaretroviral surface envelope proteins. The models are consistent with much of previously described experimental data, show regions likely to interact with receptors and identify domains that may be involved in mechanisms of antibody neutralization resistance in the small ruminant lentiviruses. The models will allow more precise design of mutants to further determine mechanisms of viral entry and immune evasion in this group of viruses and constructs for structure of these surface envelope proteins.

## Main text

The surface envelope glycoprotein of retroviruses mediate virus entry by binding to host cell receptors and are a major target of neutralizing antibodies. The surface envelope glycoprotein gp120 of the human immunodeficiency virus type 1 (HIV-1) has been structurally and functionally characterized in detail (1, 2). The surface envelope glycoproteins of other lentiviruses and the more distantly related betaretroviruses have not been characterized structurally in detail and functional information regarding receptor binding sites (RBS) is more limited in most cases due in part to the lack of structural information to guide such studies. Structural determination for the surface envelope glycoprotein of lentiviruses is particularly challenging due to the high glycosylation and, in analogy to what has been observed with HIV-1 gp120, a possibly inherent domain flexibility linked to mechanisms of receptor recognition and immune evasion (1). An additional challenge for the small ruminant caprine arthritis-encephalitis (CAEV) and visna viruses is that host cell receptors used for entry are not known.

The more salient structural aspects of HIV-1 gp120 are a two-domain organization with a four-strand “bridging sheet” minidomain emanating from both domains (1, 2). The inner domain is composed of three different layers and interacts with the transmembrane glycoprotein in the axis of the trimeric envelope structure (2). The CD4 receptor binds between the two main domains and the bridging sheet. The CD4-binding site is flanked on one side by the V1/V2 variable loop emanating from the bridging sheet in the inner domain and by the V3 loop emanating from the outer domain on the other side. These loops are important components of the mechanisms HIV-1 immune evasion. The coreceptor binding site maps to the bridging sheet formed or stabilized upon gp120 binding to CD4 (1). The outer domain of gp120 is heavily glycosylated, shielding it from antibody binding (1).

Sequence similarity is readily observed in the transmembrane glycoprotein of lentiviruses and the more distantly related betaretroviruses (3). However, the sequences of the surface envelope glycoproteins of lentiviruses and betaretroviruses is highly diverse. The sequence diversity is a consequence of the rapid sequence change, deep divergence and adaptation to different hosts. Only limited, fragmentary sequence similarity has been observed in the surface glycoproteins of lentiviruses, similarity that also extends to the betaretroviruses (4, 5, 6). This fragmentary sequence similarity is only clear in the context of the HIV-1 gp120 structure. The more conserved modules are confined to the β-sandwich of the inner proximal domain that interacts with the transmembrane region in gp120. In CAEV gp135, the inner domain model derived from these similarities was supported by results from mutagenesis studies (7). In addition, the pattern of cysteine residue and glycosylation site distribution and the location of sequence similarity modules has indicated additional structural aspects shared among lentiviral surface glycoproteins, including homologues of the gp120 V1/V2 loops in some lentiviruses and an overall two-domain organization in other lentiviruses (6, 8). More detailed information about receptor binding sites is not obvious from these models due to the lack of any additional sequence similarity in the envelope glycoprotein of lentiviruses outside the inner proximal domain. The recent release of the AlphaFold package for structural prediction (9, 10) provides an opportunity to make inroads into the structural characterization of the surface glycoprotein of these and other retroviruses to understand mechanisms of viral entry and immune evasion.

AlphaFold is an artificial intelligence computational method for structural prediction (9). The AlphaFold method has been shown to provide models that in many cases are equivalent to experimentally derived structures (9). Importantly, AlphaFold provides validated metrics for model reliability in its output, facilitating model accuracy interpretation. The mature region of the surface envelope glycoproteins of non-primate lentiviruses and betaretroviruses were modeled using the Colab server implementing a simplified AlphaFold method (https://colab.research.google.com/github/deepmind/alphafold/blob/main/notebooks/AlphaFold.ipynb). Models were analyzed with PyMOL version 2.5.1 (Schrödinger, LLC).

Modeling of the surface envelope glycoproteins of feline immunodeficiency virus (FIV), equine infectious anemia virus (EIAV), bovine immunodeficiency (BIV) and jembrana disease (JDV) lentiviruses and the consensus rabbit endogenous lentivirus RELIK yielded low reliability models with predicted Local Distance Difference Test (pLDDT) confidence scores (range, 0 to 100) of less than 70, a cut-off for main chain prediction reliability (9), and often less than 50 for most or all of the sequence (Supplementary Fig. S1). This was observed even when sequences predicted to be analogous to the gp120 V1/V2 loops were removed from the input to attempt to improve modeling performance (not shown). These models were not further considered. In contrast, high quality models were obtained for the full-length mature CAEV and visna virus gp135. The pLDDT scores for gp135 models were greater than 70 for 64% to 78% and greater than 90 for 4% to 15% of the sequences (Supplementary Fig. S1 and S2). This indicates high reliability for most of the main chain (pLDDT ≥ 70) and side-chains (pLDDT ≥ 90) of a significant fraction of the modeled residues (9). It is not known if the models reflect unliganded or receptor bound gp135 conformations. The description of the models described below follow the naming and orientation conventions used for HIV-1 gp120, with inner and outer domains and proximal and distal sections named relative to the amino and carboxy termini of the proteins and shown with the side presumed to face the host cell down (1, 2). In addition, the sides of gp135 equivalent and opposite to the CD4 binding side of HIV-1 gp120 are referred here as the R+ and R-sides, respectively.

The CAEV and visna virus gp135 models are essentially identical despite a sequence identity of 65% between the two groups of viruses. The root mean square deviation (rmsd) of the models was 1.03 to 1.66 Å including the more variable terminal regions. Therefore, both models are described in combination using the higher scoring CAEV gp135 model for illustration and highlighting key differences in the visna virus gp135 model. Extensions at the amino and carboxy termini of the protein have low pLDDT scores (Supplementary Fig. S1 and S2) and are probably not modeled in the correct position that is likely to be dependent on transmembrane subunit interactions. Those interactions are not modeled by AlphaFold (9). In fact, modeling of the entire envelope glycoprotein including the transmembrane subunit did not result in plausible models of these gp135 terminal extensions and the transmembrane subunit (not shown). These terminal regions are excluded from the analyses. All cysteine residues participate in 11 disulfide bonds (Supplementary Fig. S3). The gp135 models have a multi-domain distribution similar to HIV-1 gp120 (1, 2). However, gp135 is divided into three domains in the models rather than two as in gp120: an inner domain and two outer domains emanating from the inner domain in the same direction as the HIV-1 gp120 outer domain (Fig. 1).

**Figure 1.**
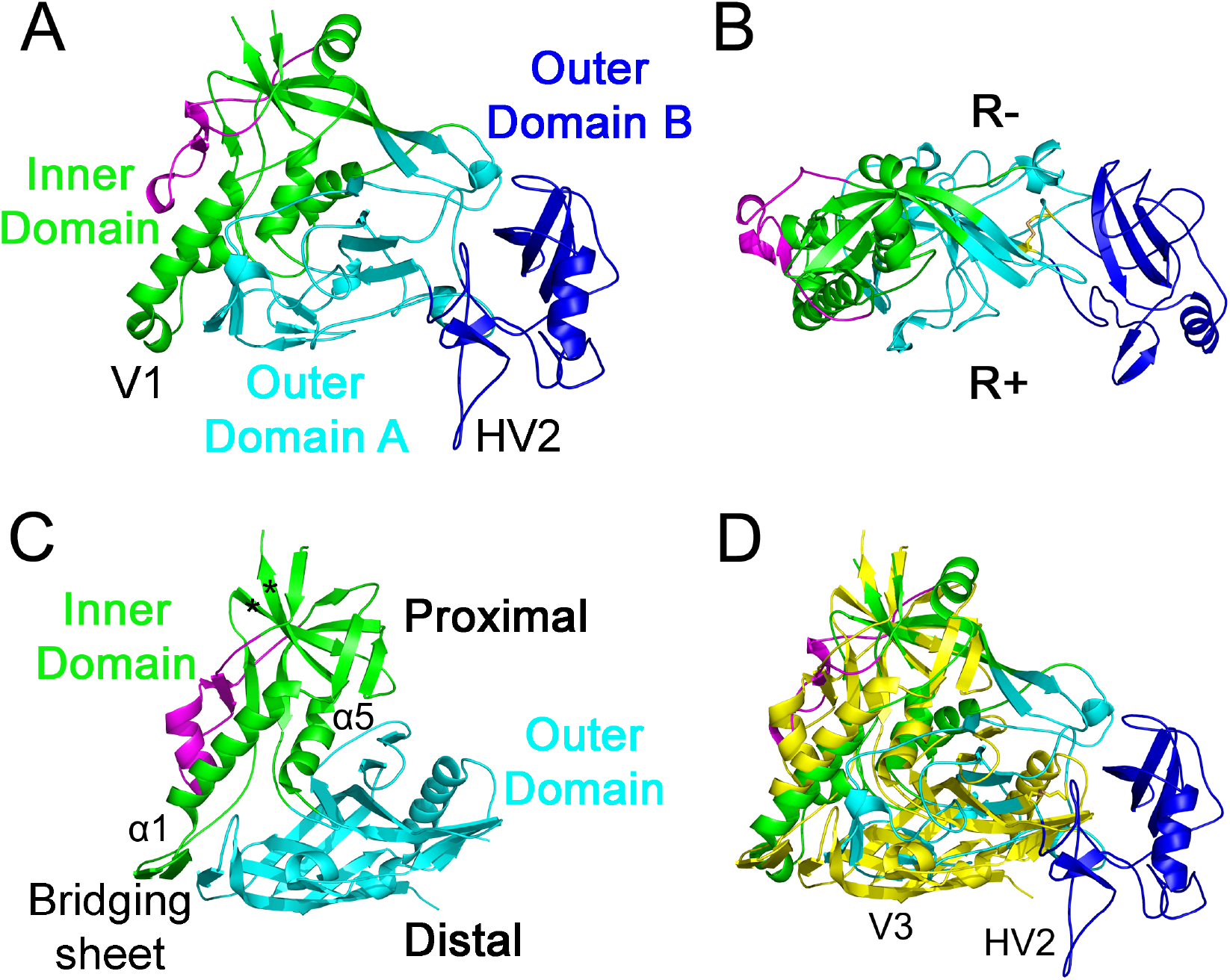
Domain organization of CAEV and visna virus gp135. Models of CAEV-63 g135 observed from (A) the R+ side and (B) from the proximal region are shown. (C) HIV-1 gp120 structure from PDB 3JWD. (D) Superpositions of the gp135 model from panel A and the gp120 structure from panel C shown in yellow. In panel C, the inner domain section within the gp120 “core” structure including layers 2 and 3 are shown in green, the inner-domain layer 1 is shown in magenta, and the outer domain in cyan. CAEV domains homologous to gp120 domains are shown in the same color scheme in panels A, B and D, showing the additional outer domain B in blue. The gp135 model in panels A and D and gp120 structures in panel C and D are aligned by the β-strands 5 and 25 of the proximal region of the gp120 inner domain (asterisks). The R+ and R-sides of gp135 are indicated in panel B. The approximate position of the gp120 V3 loop and the gp135 HV2 loop are shown in panel D. The proximal and distal regions of gp120 and the position of the bridging in CD4-bound gp120 is indicated in C. The positions of the CAEV V1 and the hypervariable HV2 region are indicated in A. The disulfide bond at the base of outer domain B is shown as sticks in shades of yellow in panel B. The GenBank accession number for CAEV-63 gp135 used for modeling is M60855. The terminal extensions are omitted for clarity.

The inner domain has an overall subdomain distribution similar to the gp120 inner domain with three layers, one outside the “core” structure and two within (2). The second and third layers of the gp135 inner domain that correspond to the gp120 inner domain regions in the “core” protein have the same elements as gp120, including the proximal β-sandwich and two helices corresponding to gp120 helices α1 and α5 (Fig. 1A and C). The first layer of the gp135 inner domain is in a position similar to the corresponding layer of gp120. The end of gp135 helix α1 is not followed by a β-sheet at the base of a long loop as in gp120 but rather by a short helical section with relatively low pLDDT scores that include the variable gp135 V1 region (Fig. 1A and B, Supplementary Fig. S2). A significant difference between gp135 and gp120 is that in gp135 a 15-residue sequence extends as a β-hairpin from the inner domain across the proximal region of the first outer domain (Fig. 1A and B). Residues in that extension have a relatively high average pLDDT score of 87.8 (Supplementary Fig. S1 and S2). The inner domain has four to five glycosylation sites, none in the likely transmembrane subunit contact region (Supplementary Fig. S4). The closest structural homolog for the gp135 inner domain model determined with the Dali server (ekhidna2.biocenter.helsinki.fi) is the clade A/E HIV-1 93TH057 strain gp120 core structure (PDB 4JB9), with an rmsd of 3 Å.

The outer domain closest to the inner domain occupies approximately the same space as the gp120 outer domain and is referred to as outer-domain A (Fig. 1). Outer domain A includes a total of 170 to 175 residues and is centered around a four-strand β-sheet, with another β-hairpin close to a homologue of the gp120 bridging sheet in the distal region of the domain A close to the inner domain (Fig. 1A and C). The distal region of outer domain A has the lowest pLDDT scores in the domain (Supplementary Fig. S1 and S2). Outer domain A includes four to nine N-linked glycosylation sites, mostly on the R-side of gp135 (Supplementary Fig. S4). No significant outer domain A structural homologs were identified except for the partial bridging sheet homolog.

Outer domain B emanates from outer domain A (Fig. 1A) and is delimited by a disulfide bond at its base in outer domain A (Fig. 1B, Supplementary Fig. S3). Outer domain B includes 137 to 139 residues and extends beyond the space corresponding to the gp120 V5 region, slightly tilting distally and towards the R+ side in the models (Fig. 1A and B). The CAEV gp135 hypervariable region HV2 that is presumed to be involved in neutralization escape during persistent infection (11) forms a loop emanating from the distal end of outer-domain B, displaced outwards relative to the gp120 V3 variable loop position (Fig. 1D). In visna virus the HV2 gp135 region has been shown to be involved in neutralization escape and is named the principal neutralization domain (12, 13). This region in visna virus gp135 includes deletions in some strains relative to CAEV gp135, which results in a shorter loop within outer domain B (Supplementary Fig. S5). Outer domain B includes nine to ten N-linked glycosylation sites, covering much of the domain except the R+ side (Supplementary Fig. S4). Despite the overall size similarity of gp120 and gp135, the sequences have significantly different arrangements in both proteins. Whereas in gp120 a significant fraction of the sequence is included in the long V1/V2 and V3 variable loops emanating from the inner and outer domains (1), gp135 has only one short variable loop (Fig. 1A), with much of the balance of the sequence forming an additional outer domain folding independently. Whether the primate and the small ruminant lentiviruses descend from a common ancestor with a surface envelope glycoprotein with one or two outer domains is not clear. No significant outer domain B structural homologs were identified.

The interface of the two outer domains form a deep cleft on the R+ side of the gp135 model (Fig. 1B and 2B and D). This cleft is located in a position analogous to the CD4 binding site of gp120 (1) but displaced slightly outwards relative to the CD4 binding site (Fig. 2). The surfaces bordering that cleft may constitute a gp135 RBS. Supporting this interpretation, the cleft is devoid of glycosylation sites, is flanked by sequences more conserved among disparate CAEV and visna virus strains in outer domain A and is proximal to the bridging sheet β-hairpin homologue on one side and the hypervariable loop on the other (Fig. 1A and B, Supplementary Fig. S4). The CD4 binding site of gp120 shares the same properties except that it is located mostly in the outer domain with some contribution from the inner domain (1), rather than between two outer domains as in the gp135 model (Fig. 2A). The putative gp135 RBS in the outer domain cleft is lined with several conserved residues on the outer domain A side (Supplementary Fig. S3 and S4). The same surface includes a positive charge patch of variable size in different strains (Supplementary Fig. S6). Residues that are divergent between CAEV and visna virus strains cluster on the R-side and the distal region of the outer domains, together with glycosylation sites (Supplementary Fig. S4). These regions are less likely to include a gp135 RBS due to these sequence properties. Flexibility between the two gp135 outer domains hinging around the disulfide-constrained domain link is a possibility, which may lead to modulation of RBS accessibility. One unexpected finding is that the gp135 region analogous to the bridging sheet homologue in outer domain A and the distal helical region of the inner domain is composed of residues that are variable between visna virus and CAEV (Supplementary Fig. S3 and S4). Whether this reflect the differential receptor usage of these viruses (14, 15) or whether these regions do not participate in receptor or coreceptor binding is not clear.

**Figure 2.**
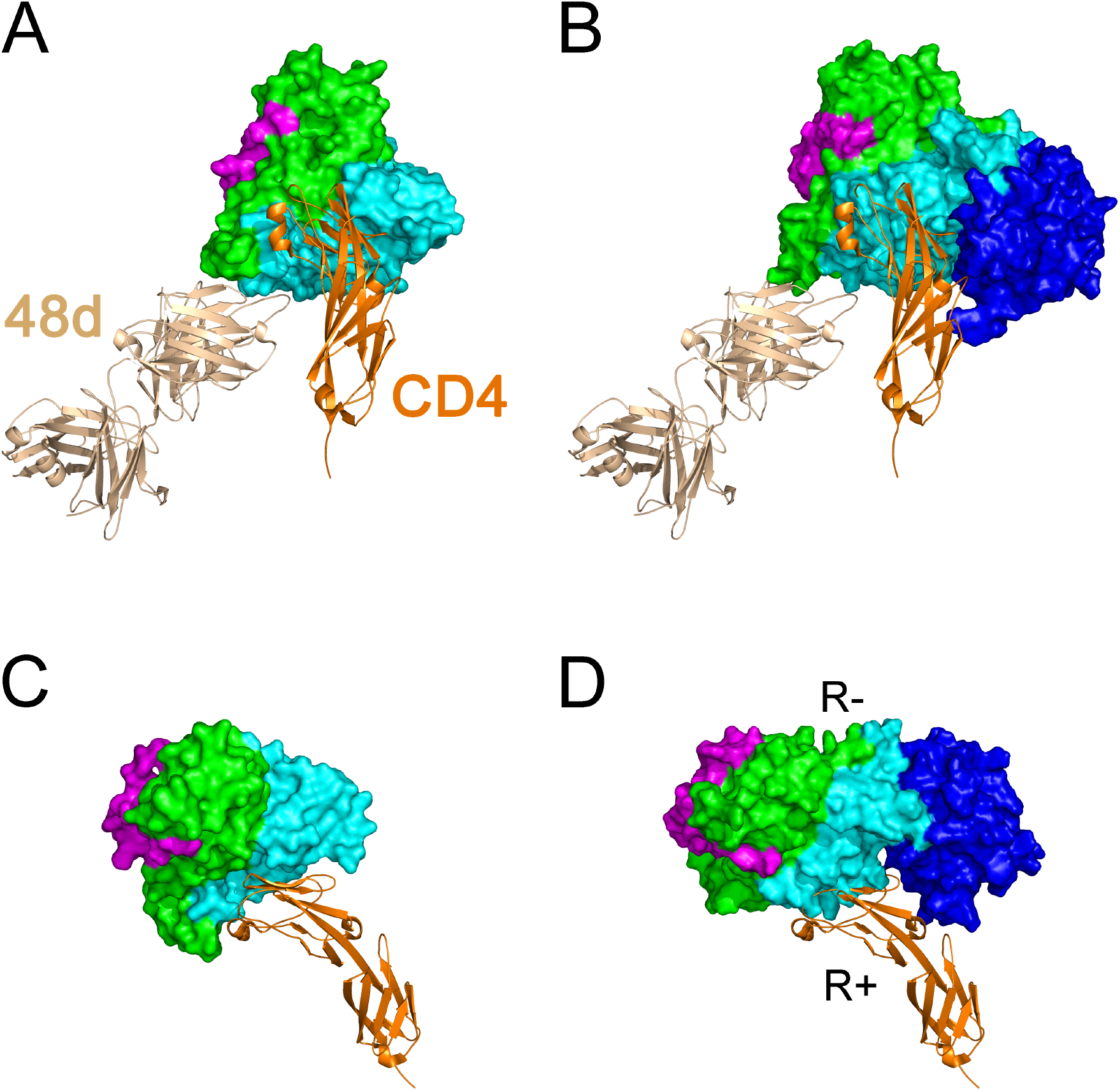
Location of the putative CAEV gp135 RBS. (A) Structure of HIV-1 gp120 in complex with CD4 and antibody 48d binding an epitope overlapping the gp120 coreceptor binding site (PDB 3JWD). (B) CAEV gp135 in the same orientation showing the approximate regions of gp135 that correspond to the CD4 and antibody 48d bindings sites of gp120. Panels C and D show the same structure and model shown in A and B from the proximal (top in A and B) side, omitting antibody 48d for clarity. Domain colors are the same as for Figure 1 in all panels. The HIV-1 gp120 structure and CAEV gp135 model are aligned by the HIV-1 gp120 β-strands 5 and 25 as in Figure 1. Amino and carboxy terminal extensions are omitted for clarity.

Previous data supports several aspects of the gp135 models. As mentioned earlier, mutagenesis studies of the proximal region of the CAEV gp135 inner domain are consistent with this region interacting with the transmembrane glycoprotein (7). The location of the CAEV gp135 hypervariable loop HV2 and the visna virus gp135 principal neutralization domain nearby the putative RBS is consistent with a role in antibody neutralization escape. Non-neutralizing goat antibody F7-299, which cross-reacts with CAEV and visna virus gp135 (16), binds an epitope overlapping a gp135 RBS and includes two large negative charge patches in its complementarity-determining regions (Supplementary Fig. S7). A likely location for the F7-299 epitope overlapping an RBS is thus the positively charged and relatively conserved region in outer domain A overlapping the putative RBS, as the rest of the outer domain surface is otherwise covered with variable residues or glycosylation sites. Mutation of CAEV gp135 Ile-166 to Alanine results in increased fusogenic activity, increased binding of RBS-specific antibody F7-299 to soluble and oligomeric CAEV gp135 and increased conformational stability or accessibility of the F7-299 epitope (16). The model shows Ile-166 in the inner domain forming a hydrophobic interaction with Ile-304 and Ile-306 in the bridging sheet homologue in the distal region of outer domain A (Supplementary Fig. S8). Disruption of that interaction may allow the β-hairpin to move or may allow conformational changes involving that region and the adjacent putative RBS. This would be analogous to the coordinated conformational changes of the CD4 binding site and the coreceptor binding site in the bridging sheet of gp120 (1). Finally, deletion of the variable gp135 V1 region within the distal helical region of the inner domain results in increased cell-to-cell fusion and binding by RBS-specific antibody F7-299 to the trimeric, but not soluble monomeric, envelope glycoprotein (16, 17), indicating that this region may participate in RBS-occluding interactions with other subunits of the trimer in a manner analogous to the similarly located gp120 V1/V2 loop (18).

Models of the surface envelope proteins of the betaretroviruses jaagsiekte sheep retrovirus (JSRV), mouse mammary tumor virus (MMTV) and consensus human endogenous retrovirus type K (HERV-K) had high pLDDT scores of 70 or above for 79% to 89% and 90 or above for 30% to 54% of the sequence (Supplementary Fig. S1). The betaretrovirus surface envelope protein models have a single domain. The amino and carboxy terminal ends of betaretroviral envelope proteins also extend away from the inner domain homologues and are probably not well-modeled (not shown). Four disulfide bonds are modeled in the same locations for JSRV and MMTV surface envelope proteins whereas in the consensus HERV-K model only 4 of the 8 cysteine residues form disulfide bonds (Supplementary Fig. S9 and S10). Whether the unpaired cysteines point to inconsistencies in the HERV-K model, reflect ambiguities from consensus sequence derivation (19) or both is not clear.

No clear structural similarity was identified between the betaretroviral surface envelope protein models shown here and the receptor domain structures of murine and feline leukemia virus gammaretroviruses (20, 21). Instead, as expected from the previously identified sequence similarities in the surface envelope and transmembrane subunits of lentiviruses and betaretroviruses and from the phylogenetic clustering of betaretroviruses and lentiviruses (3, 5, 6, 22), the surface envelope proteins of betaretroviruses share structural elements with HIV-1 gp120. The betaretroviral models show a β-sandwich structure that is highly conserved among the three betaretroviruses and similar to the same inner-proximal domain β-sandwich of HIV-1 gp120 (Fig. 3, Supplementary Fig. S11). The betaretroviral envelope protein models lack the helical regions that are observed in the gp120 inner domain. Using the gp120 β-strands 5 and 25 homologues as reference points to orient the models, the betaretroviral surface envelope proteins also have a region equivalent to the gp120 inner-domain layer 1 but no outer domain. The region topologically equivalent to the outer domain in the betaretroviruses, between the gp120 β-strands 8 and 25 homologues, is short and confined to the proximal region (Supplementary Fig. S9 and S11). Instead, the betaretroviruses have long insertion relative to gp120 in a region topologically equivalent to gp120 helix α1, the first part of the bridging sheet and the loop connecting back to inner domain β-strand 4, but otherwise structurally distinct from gp120 (Supplementary Fig. S11). This insertion makes up a mid-domain region fairly conserved structurally among the betaretroviruses but distinct form the lentiviruses and apical regions directed away from the virion surface that are more structurally diverse among the betaretroviruses with varying lengths for the four apical excursions in each virus (Fig. 3). These apical regions are likely candidates for the betaretroviral RBS. The observed structural differences in the apical regions of these surface envelope proteins may thus reflect differences in receptor specificity.

**Figure 3.**
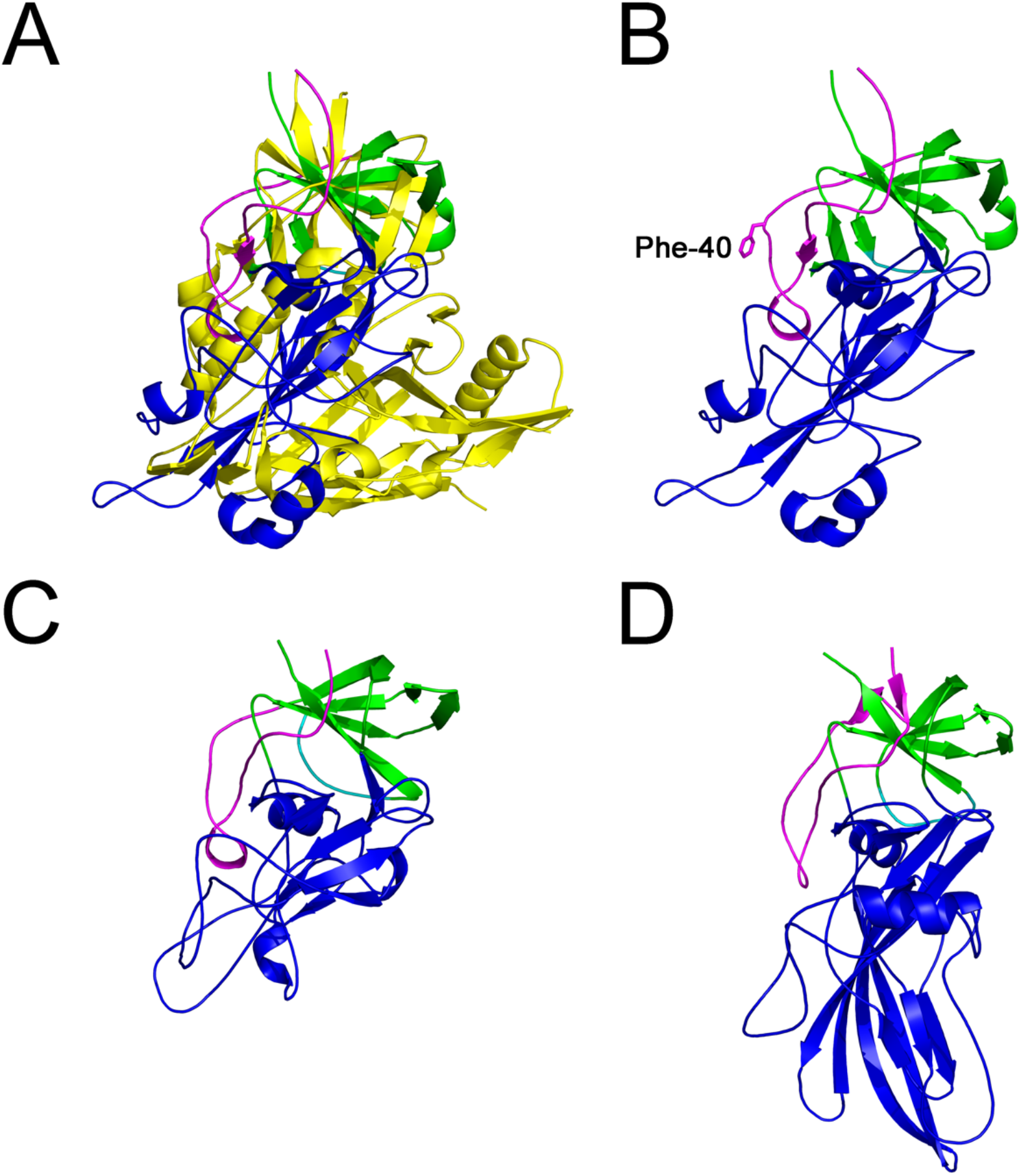
Betaretrovirus surface envelope protein structural models. (A) Superposition of the HIV-1 gp120 structure (yellow) and MMTV surface envelope protein showing no overlap in the HIV-1 gp120 outer-domain region. Structural models of (B) MMTV (C) JSRV and (C) consensus HERV-K envelope proteins showing the layer 1 inner domain homologue in magenta, the inner-proximal domain homologue in green and the apical regions in blue. All models are shown in the same orientation aligned by the conserved β-strand 5 and 25 homologues. The Phe-40 residue of the MMTV virus surface envelope protein is shown in panel B. Genbank accession numbers for the sequences of the models in panels B and C are M80216 and X01811. The model in panel D is based on a consensus HERV-K surface envelope glycoprotein sequence (19).

It has been suggested that the region around residue 40 of the MMTV surface envelope protein interacts with its receptor, mouse transferrin receptor (23, 24). However, in the MMTV model the region around residue 40 faces the expected trimer interface in the proximal region of the protein, which would make it unavailable for receptor binding (Fig. 3B). The effects of residue 40 mutations on increased receptor binding activity that have been described (23) may thus be indirect to perturbations in inter-subunit interactions, similar to inner domain mutations in CAEV gp135 that expand host range and increase envelope glycoprotein fusion activity (7). Further functional studies based on the models provided here should resolve this apparent discrepancy.

The models described here point to evolutionary relationships and divergences between the surface envelope proteins of divergent retroviral groups. The apical sequences with the potential betaretroviral RBS regions are topologically equivalent to the distal region of the gp120 inner domain that forms the bridging sheet involved in coreceptor binding, between the strand β1 and β4 homologues (Supplementary Fig. S11). This is in contrast with HIV-1 and other lentiviruses, in which RBS regions and coreceptor binding sites are also included in a long, multi-domain, sequence between major elements of the inner-proximal domain. The region topologically equivalent to the outer domain in the betaretroviruses is short and unavailable for receptor binding. This points to the evolution of equivalent and non-equivalent structures for host cell interactions in the surface envelope proteins of lentiviruses and betaretroviruses based on the same conserved inner-proximal domain framework.

The AlphaFold algorithm provides high quality models of CAEV and visna virus gp135 and betaretroviral envelope glycoproteins that are self-consistent and supported by existing data with the possible exception of the MMTV RBS. Most CAEV and visna virus strains are resistant to antibody neutralization (25-28). The structural models point to factors that differ structurally from those mediating neutralization resistance in HIV-1, including the absence of long variable loops and the presence of an additional outer domain that may participate in RBS occlusion. The models presented here should guide further structural and functional characterization of these envelope glycoproteins, enabling the precise design of point and deletion mutants to further dissect the mechanisms of viral entry and immune evasion in these viruses and possibly engineering altered receptor specificity and cell tropism of betaretroviral envelope proteins for therapeutic applications.

## Acknowledgments

The author thanks Seth Harris for help running AlphaFold and Peter Hsu with structural homolog searches. PDB files with the gp135 and betaretroviral surface envelope protein model coordinates are available upon request.

## References

1. Kwong, P.D., R. Wyatt, J. Robinson, R.W. Sweet, J. Sodroski, and W.A. Hendrickson. 1998. Structure of an HIV gp120 envelope glycoprotein in complex with the CD4 receptor and a neutralizing human antibody. Nature 393:648–659.

2. Pancera, M., S. Majeed, Y.A. Ban, L. Chen, C. Huang, L. Kong, Y.D. Kwon, J. Stuckey, T. Zhou, J.E. Robinson, W.R. Schief, J. Sodroski, R. Wyatt, and P.D. Kwong. 2010. Structure of HIV-1 gp120 with gp41-interactive region reveals layered envelope architecture and basis of conformational mobility. Proc. Natl. Acad. Sci. USA 107:1166–1171.

3. Schulz, T.F., B.A. Jameson, L. Lopalco, A.G. Siccardi, R.A. Weiss, J.P. Moore. 1992. Conserved structural features in the interaction between retroviral surface and transmembrane glycoproteins? AIDS Res. Hum. Retroviruses 8:1571–1580.

4. Hötzel, I., and W.P. Cheevers. 2000. Sequence similarity between the envelope surface unit (SU) glycoproteins of primate and small ruminant lentiviruses. Virus Res. 69:47–54.

5. Hötzel, I., and W.P. Cheevers. 2001. Conservation of human immunodeficiency virus type 1 gp120 inner-domain sequences in lentivirus and type A and B retrovirus envelope surface glycoproteins. J. Virol. 75:2014–2018.

6. Hötzel, I. 2008. Conservation of inner domain modules in the surface envelope glycoproteins of an ancient rabbit lentivirus and extant lentiviruses and betaretroviruses. Virology 372:201–207.

7. Hötzel, I., and W.P. Cheevers. 2003. Caprine arthritis-encephalitis virus envelope surface glycoprotein regions interacting with the transmembrane glycoprotein: structural and functional parallels with human immunodeficiency virus type 1 gp120. J. Virol. 77:11578–11587.

8. Hötzel, I., 2003. Conservation of the human immunodeficiency virus type 1 gp120 V1/V2 stem/loop structure in the equine infectious anemia virus (EIAV) gp90. AIDS Res. Hum. Retroviruses 19:923–924.

9. Jumper J., R. Evans, A. Pritzel, T. Green, M. Figurnov, O. Ronneberger, K. Tunyasuvunakool, R. Bates, A. Žídek, A. Potapenko, A. Bridgland, C. Meyer, S.A.A. Kohl, A.J. Ballard, A. Cowie, B. Romera-Paredes, S. Nikolov, R. Jain, J. Adler, T. Back, S. Petersen, D. Reiman, E. Clancy, M. Zielinski, M. Steinegger, M. Pacholska, T. Berghammer, S. Bodenstein, D. Silver, O. Vinyals, A.W. Senior, K. Kavukcuoglu, P. Kohli, and D. Hassabis. 2021. Highly accurate protein structure prediction with AlphaFold. Nature https://doi.org/10.1038/s41586-021-03819-2.

10. Tunyasuvunakool, K., J. Adler, Z. Wu, T. Green, M. Zielinski, A. Žídek, A. Bridgland, A. Cowie, C. Meyer, A. Laydon, S. Velankar, G.J. Kleywegt, A. Bateman, R. Evans, A. Pritzel, M. Figurnov, O. Ronneberger, R. Bates, S.A.A. Kohl, A. Potapenko, A.J. Ballard, B. Romera-Paredes, S. Nikolov, R. Jain, E. Clancy, D. Reiman, S. Petersen, A.W. Senior, K. Kavukcuoglu, E. Birney, P. Kohli, J. Jumper, and D. Hassabis. 2021. Highly accurate protein structure prediction for the human proteome. Nature https://doi.org/10.1038/s41586-021-03828-1.

11. Hötzel, I., N. Kumpula-McWhirter, and W.P. Cheevers. 2002. Rapid evolution of two discrete regions of the caprine arthritis-encephalitis virus envelope surface glycoprotein during persistent infection. Virus Res. 84:17–25.

12. Skraban, R., S. Matthíasdóttir, S. Torsteinsdóttir, G. Agnarsdóttir, B. Gudmundsson, G. Georgsson, R.H. Meloen, O.S. Andrésson, K.A. Staskus, H. Thormar, and V. Andrésdóttir. 1999. Naturally Occurring Mutations within 39 Amino Acids in the Envelope Glycoprotein of Maedi-Visna Virus Alter the Neutralization Phenotype. J. Virol. 73:8064–8072.

13. Haflidadóttir, B.S., S. Matthíasdóttir, G. Agnarsdóttir, S. Torsteinsdóttir, G. Pétursson, O.S. Andrésson, and V. Andrésdóttir. 2008. Mutational analysis of a principal neutralization domain of visna/maedi virus envelope glycoprotein. J. Gen. Virol. 89:716–721.

14. Jolly, P.E., and O. Narayan. 1989. Evidence for interference, coinfections, and intertypic virus enhancement of infection by ovine-caprine lentiviruses. J. Virol. 63:4682–4688.

15. Hötzel, I., and W.P. Cheevers. 2001. Differential receptor usage of small ruminant lentiviruses in ovine and caprine cells: host range but not cytopathic phenotype is determined by receptor usage. Virology 301:21–31.

16. Hötzel, I., and W.P. Cheevers. 2005. Mutations increasing exposure of a receptor binding site epitope in the soluble and oligomeric forms of the caprine arthritis-encephalitis lentivirus envelope glycoprotein. Virology 339:261–272.

17. Valas, S., C. Benoit, C. Baudry, G. Perrin, and R.Z. Mamoun. 2000. Variability and immunogenicity of caprine arthritis-encephalitis virus surface glycoprotein. J. Virol. 74:6178–6185.

18. Rusert, P., A. Krarup, C. Magnus, O.F. Brandenberg, J. Weber, A.K. Ehlert, R.R. Regoes, H.F. Günthard, and A. Trkola. 2011. Interaction of the gp120 V1V2 loop with a neighboring gp120 unit shields the HIV envelope trimer against cross-neutralizing antibodies. J. Exp. Med. 208:1419–1433.

19. Dewannieux, M., F. Harper, A. Richaud, C. Letzelter, D. Ribet, G. Pierron, and T. Heidmann. 2006. Identification of an infectious progenitor for the multiple-copy HERV-K human endogenous retroelements. Genome Res. 16, 1548–155.

20. Fass, D., R.A. Davey, C.A. Hamson, P.S. Kim, J.M. Cunningham, and J.M. Berger. 1997. Structure of a murine leukemia virus receptor-binding glycoprotein at 2.0 angstrom resolution. Science 277:1662–1666.

21. Barnett, A.L., D.L. Wensel, W. Li, D.F. Cunningham, and J.M. Cunningham. 2020. Structure and mechanism of a coreceptor for infection by a pathogenic feline retrovirus. J. Virol. 77:2717–2729.

22. Gifford, R. 2012. Viral evolution in deep time: lentiviruses and mammals. Trends Genet. 28:89–100.

23. Konstantoulas C.J., B. Lamp, T.H. Rumenapf, and S. Indik. 2015. Single amino acid substitution (G42E) in the receptor binding domain of mouse mammary tumour virus envelope protein facilitates infection of non-murine cells in a transferrin receptor 1-independent manner. Retrovirology 12:43.

24. Zhang, Y., J.C. Rassa, M.E. deObaldia, L.M. Albritton, and S.R. Ross. 2020. Identification of the receptor binding domain of the mouse mammary tumor virus envelope protein. J. Virol. 77: 10468–10478.

25. Cheevers, W.P., T.C. McGuire, L.K. Norton, R. Cordery-Cotter, and D.P. Knowles. 1993. Failure of neutralizing antibody to regulate CAE lentivirus expression in vivo. Virology 196:835–839.

26. Cheevers, W.P., D.P. Knowles Jr., and L.K. Norton. 1991. Neutralization-resistant antigenic variants of caprine arthritis-encephalitis lentivirus associated with progressive arthritis. J. Infect. Dis. 164:679–685.

27. Narayan, O., D. Sheffer, D.E. Griffin, J. Clements, and J. Hess. 1984. Lack of neutralizing antibodies to caprine arthritis-encephalitis lentivirus in persistently infected goats can be overcome by immunization with inactivated *Mycobacterium tuberculosis*. J. Virol. 49:349–355.

28. Thormar, H., M.R. Barshatzky, K. Arnesen, and P.B. Kozlowski. 1983. The emergence of antigenic variants is a rare event in long-term visna virus infection in vivo. J. Gen. Virol. 64:1427–1432.

